# Longitudinal analysis of the hand microbiome in response to chlorine-based antiseptic use during a military field exercise

**DOI:** 10.1101/2025.11.20.689467

**Authors:** Susanne Glenna, Einar E. Birkeland, Russell J. S. Orr, Gregor D. Gilfillan, Marianne Dalland, Ole Andreas Økstad, Øyvind A. Voie, Trine B. Rounge

## Abstract

Hand hygiene is essential for infection control, yet the impact of frequent antiseptic use on the skin microbiome, which is crucial for skin barrier function and pathogen exclusion, remains underexplored, especially in field conditions. In high-risk military settings, there is a need for safe, multipurpose antiseptics that avoid the drawbacks of alcohol-based options, including skin irritation, flammability, and unsuitability for wound application. We conducted a longitudinal study during the Norwegian-led military exercise *Cold Response 2022* to assess the effects of repeated use of stabilized hypochlorous acid, a non-alcoholic antiseptic, on the hand microbiome of soldiers in a field environment. Participants used either the chlorine-based antiseptic (*n* = 20) or standard hygiene practices, mainly alcohol-based sanitizers (*n* = 19), for 10 days. Skin swabs from hands and untreated forearms were collected at baseline, post-intervention, and three weeks after discontinuation, and analyzed using 16S rRNA sequencing. Overall, our data revealed that field exposure accounted for more variation in bacterial composition than antiseptic use. When comparing alpha and beta diversity trajectories between the two antiseptic groups, there was minimal divergence during active field use, but a significant post-treatment increase in alpha diversity associated with the use of chlorine-based antiseptic. In contrast, untreated forearms showed no such group-level differences. These findings suggest that antiseptic-driven microbiome shifts are most apparent during recovery, with non-alcoholic formulations supporting faster recolonization after treatment. Stabilized hypochlorous acid represents a promising alternative to standard alcohol-based antiseptics for military personnel.

**IMPORTANCE:** Effective hand hygiene reduces pathogen transmission, but its influence on the skin microbiome has rarely been studied. Our research emphasizes the importance of longitudinal assessments by identifying greater antiseptic-driven differences occurring after cessation of treatment than during repeated use. Service members frequently use antiseptics in challenging environments, highlighting the operational need for a non-alcoholic, multipurpose product that can be safely used on both intact skin and wounds. Our findings suggest that a chlorine-based non-alcoholic antiseptic permits faster recovery of bacterial diversity after perturbation, informing product selection and hygiene policies in defense and other high-risk settings.

## INTRODUCTION

The skin acts as a protective barrier against environmental threats, with its microbiota - comprising both resident and transient microorganisms - playing a central role in maintaining cutaneous homeostasis and preventing pathogen colonization (1–3). Among all skin sites, the hands are uniquely exposed to external influences, making their microbial communities highly dynamic and sensitive to factors such as hygiene practices (2, 4, 5). Hand hygiene is one of the most effective measures for reducing pathogen transmission (6–9), yet the effects of hand washing and sanitizer use on microbial diversity and resilience in different populations remain poorly understood (10–13). This limits our ability to balance hygiene with microbiome preservation. Most antiseptics, such as chlorhexidine and povidone-iodine, are used in a broad-spectrum, non-selective manner, reducing microbial burden indiscriminately across taxa (14, 15). While effective for infection control, such non-targeted action can also disrupt commensal as well as pathogenic populations and alter community composition, with recovery patterns varying by agent and context (10, 11, 16). As the global use of topical antimicrobial agents continues to rise (17), longitudinal studies are needed to evaluate their effects on a microbial community-wide scale (18, 19).

In military field conditions, where infection and injury risks are elevated, good personal hygiene and effective antimicrobial agents are critical (20–24). Military personnel commonly use alcohol-based products for disinfecting hands and surfaces, vital for preventing infections (25, 26). However, concerns remain about their safety, as these disinfectants may cause skin irritation, especially in sensitive or compromised skin, leading to discomfort and reduced compliance (27–29). Ethanol, a common active ingredient, is irritating to wounds and mucosal surfaces, toxic if ingested, and highly flammable (30, 31). Despite these limitations, alcohol-free alternatives have not yet been successfully implemented in the defence sector. Moreover, current field protocols require multiple products for disinfection and wound care, highlighting the need for a multipurpose, alcohol-free antiseptic (25). A promising candidate is a novel combination of two well-known antimicrobial substances, hypochlorous acid (HOCl), a naturally occurring oxidant produced by our immune system (32–34), and acetic acid, historically used in medicine (35–37) and working as a buffer to stabilize HOCl at its optimal pH (32, 38). Both compounds exhibit broad-spectrum antimicrobial and anti-biofilm activity, supporting their use in treating infected wounds and skin conditions such as eczema (37, 39–44).

Despite the demonstrated *in vitro* antimicrobial activity (32, 39, 45), the influence of stabilized HOCl on the skin microbiome under real-world conditions is unknown. This study aims to explore how the field application of stabilized HOCl affects human hand bacterial populations, as part of the development of a military-grade antiseptic suitable for both intact skin and wounds. 16S rRNA amplicon sequencing was employed to assess shifts in bacterial communities in response to hand sanitation. We previously characterized the skin microbiome on hands and forearms of soldiers following a 10-day military field exercise with standard hygiene practices, showing that both skin sites exhibited temporal changes in composition, with soil- and water-associated bacteria enriched post-exercise, although hands appeared more resilient (46).

Building on these findings, the current study investigates whether HOCl-based antiseptic use during field exercise alters hand bacterial composition and diversity beyond environmental factors, through direct comparison with the previously studied control group. We compare changes between the two groups occurring during and after the repeated antiseptic intervention by evaluating samples collected before, immediately after, and following a 3-week recovery period.

## MATERIALS AND METHODS

### Study design and participants

Norwegian soldiers participating in NATO’s “*Cold Response”* exercise in Innlandet county of Norway in March 2022 were recruited for this longitudinal study. Ethical approval was obtained from the Norwegian Regional Committee for Medical and Health Research Ethics in South-Eastern Norway (ref: 359040), and all participants provided written informed consent. Participants were assigned to either a chlorine-based antiseptic intervention group or a control group following their regular hygiene practices, based on existing troop affiliations (Fig. 1). Both groups operated in the same geographical area and engaged in similar tasks during the exercise, ensuring comparable environmental exposures. The intervention group received a hypochlorous acid + acetic acid hand disinfectant (concentrations specified below) along with standardized instructions: apply ∼3 mL per use, ensure full hand coverage, allow air-drying, and apply before and after meals as well as after restroom visits over the 10-day field exercise. Each participant was issued two 100 mL flasks labeled with study IDs, weighed before and after the exercise to quantify individual usage. Participants were instructed not to share flasks and to use only the provided product. The control group received no guidance on antiseptic use and was not given access to the chlorine-based product. To avoid interfering with established routines or compromising hygiene, control group participants maintained their typical practices, which were documented. Both groups underwent identical sampling protocols at three time points (Fig. 1): 1) baseline (the day before the exercise), 2) post-intervention (after the 10-day field exercise w/wo antiseptic intervention), and 3) follow-up (three weeks after the end of exercise and antiseptic intervention). Two skin sites were sampled; the hands, which received antiseptic treatment, and the forearms, serving as untreated intra-individual controls. Exclusion criteria included self-reported antibiotic use within six months before enrollment, observable dermatologic conditions, immunocompromised states, and other self-reported illnesses. Three participants were excluded from the second and third samplings due to COVID-19 illness, and two missed the follow-up for unrelated reasons. Participants were instructed to refrain from showering, handwashing, or using skincare products for 12 hours before each sampling session. At each session, participants completed a questionnaire including questions related to their demography, health, activity, and antiseptic usage.

**Figure 1:**
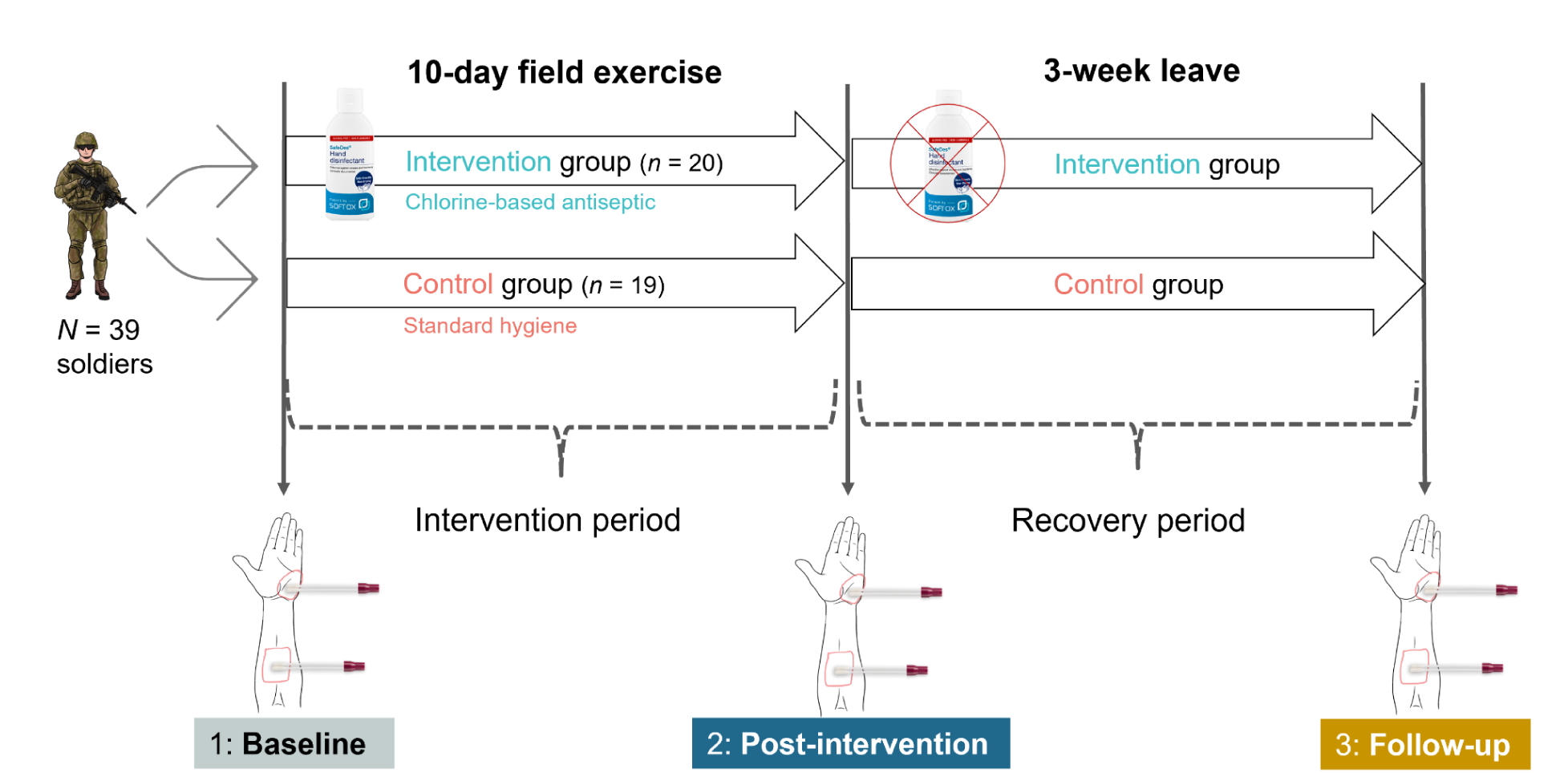
Study design. Soldiers (*n* = 39) participating in a 10-day field exercise were sampled for skin microbiome analysis at three time points: (1) baseline (pre-exercise), (2) post-intervention (immediately after exercise/treatment), and (3) follow-up (three weeks later). Participants were assigned to an intervention group (blue) using chlorine-based antiseptic or a control group (pink) maintaining standard hygiene during the exercise. Skin swabs from hands (treated) and forearms (untreated) were collected at each time point. Periods between sampling are termed intervention (exercise plus antiseptic use) and recovery (three weeks post-use).

### Antiseptic intervention product

The active-chlorine hand disinfectant (SafeDes^TM^, SoftOx Solutions AS, Norway) contains 0.02 % hypochlorous acid (HOCl) stabilized in 0.25% acetic acid (HOAc) to a pH of 4.3. It is approved according to EN 1500, EN 13727, and EN 13624.

### Sample collection

Skin samples were collected from the participants at three time points following a standardized protocol previously described (46). Briefly, skin swabs were taken from the hypothenar region of the palms and the volar forearms (Fig. 1) using sterile, pre-wetted, dual-flocked swabs. For each skin site, the left and right sides were pooled into a single sample using the same swab. Negative control swabs were included in each sampling event. All samples were immediately stored at -80°C until further processing.

### DNA extraction and 16S rRNA sequencing

The DNA extraction, library preparation, and sequencing were performed concurrently for both studies of the complete dataset, with all samples mixed randomly and processed collectively (46). In brief, total DNA was extracted from skin samples using the NucleoSpin® 96 Soil kit (Macherey-Nagel). Each swab pair was subjected to chemical (SL2 lysis buffer) and mechanical (bead-beating) lysis, after which the lysates were combined for DNA binding and purification using a spin column. Amplicon-based sequencing of the extracted DNA was performed using the Illumina protocol (47), targeting the *16S* V3-V4 region with 341F/785R primers (48). Libraries were sequenced on an Illumina MiSeq v3 with 300 bp paired-end reads.

### Bioinformatics analyses

Microbial community analyses were performed using the bioinformatics pipeline described in (46). Briefly, we performed quality-control and trimming of the 16S amplicon reads with multiQC (v1.7) (49), Trim Galore (v0.6.6) (50), and Cutadapt (v4.2) (51). Samples with <5000 reads post trimming were excluded before analysis with QIIME 2 (v2024.2) (52). The DADA2 plugin (53) was used to perform amplicon sequence variant (ASV) classification, with the SILVA 16S rRNA v138 database (54) and the q2-feature-classifier classify-sklearn (using a V3-V4 trained classifier) (55). Mitochondrial-, chloroplast-, and singleton-reads were removed before the ASV tables were imported into R for subsequent microbiome analyses using the *phyloseq* (v1.42.0) package (56).

Taxonomic classification accuracy was previously validated using mock communities processed alongside all study samples, as described in (46). The Decontam R package was used to identify and remove contaminating ASVs (Fig. S1). All samples were normalized to a sampling depth of 7000 reads based on rarefaction (Fig. S2) before statistical comparisons (unless stated otherwise).

### Statistical analysis

Statistical analyses were done in R (v4.4.1) using ggplot2 (v3.5.0) for visualizations (57). In all statistical models, the DNA extraction batch was included to adjust for technical biases, and associations considered significant at the *P* < 0.05 level. Each skin site (hands and forearms) was analyzed separately.

Alpha (within-sample) diversity was assessed using Shannon and Inverse Simpson indices. Linear mixed models were fitted using the *nlme* package (v3.1-165) to test differences in alpha diversity over time, with a random intercept for subject ID to account for repeated measures. Model assumptions were checked by inspecting residuals for normality and heteroscedasticity, and model fit was evaluated using the Akaike Information Criterion (AIC). Competing models were compared using likelihood ratio tests via ANOVA to assess whether increased complexity improved fit. To test for group-by-time interaction, we used the model: *alpha diversity ∼ group + time_point + group:time_point + batch, random= ∼1 | subject*.

Estimated marginal means (EMMs) for the fitted model were computed using the *emmeans R-*package (v1.8.9), with custom contrasts to compare the difference-in-differences between the two groups. Multiple comparisons were adjusted with the Tukey method. To test for a dose-response relationship between antiseptic use and alpha diversity after intervention, we used the following model: *post_alpha ∼ antiseptic amount + group + baseline_alpha, random = ∼1 | subject*, including baseline diversity as a covariate to adjust for initial differences.

Beta diversity was evaluated using Bray-Curtis dissimilarities. Associations were tested using PERMANOVA (999 permutations) with the *adonis2* function from the *vegan* package (v2.6-4), incorporating a group × time point interaction and subject-level stratification to account for repeated measures in the treated skin site (58). For the forearm site, which had sparse repeated measures, subject ID was included as a covariate in the model to adjust for individual-level variation. Group dispersions were assessed with PERMDISP, using *vegan*’s betadisper (type = ”centroid”) followed by permutest (999 permutations) for pairwise comparisons. When PERMDISP revealed unequal variance between groups, ANOSIM was used as a complementary test. Intra-individual Bray–Curtis changes between groups were compared using the Wilcoxon rank-sum test. A linear regression model was used to determine whether antiseptic volume (scaled) predicted changes in beta diversity within the intervention group.

Differential abundance analysis of raw ASV counts was performed with DESeq2 (59) as described earlier (46), only now with the following design formula:

*design = ∼ batch + group + time_point + group:time_point,*

where the group:*time_point* interaction identified taxa with differential trajectories in the intervention versus control groups through pairwise contrasts between time points. ASVs with an FDR-adjusted (Benjamini-Hochberg) *p*-value < 0.05 and absolute log2 fold change ≥ 1 in the interaction term were considered statistically significant. The log2 fold change reflects how the change in abundance over time differs between groups; positive values indicate that taxa increased more (or decreased less) in the intervention group compared to the control group, whereas negative values indicate that taxa increased more (or decreased more strongly) in the control group compared to the intervention group. ComplexHeatmap (v2.14.0) (60) was used to generate a heatmap of differentially abundant taxa.

## RESULTS

### Study cohort and sequencing overview

All study participants (*n* = 39) were healthy Norwegian male professional soldiers, aged 20 to 38 (median age 23), and the two groups (control, *n* = 19; intervention, *n* = 20) had comparable baseline characteristics (Table 1).

**Table 1:** Characteristics of study participants (*n* = 39) and samples in control and intervention group.

| Variables | Control ($n = 19$) | Intervention ($n = 20$) |
| --- | --- | --- |
| Age<sup>a</sup> | 23.6 ± 3.1 (20-30) | 24.7 ± 4.2 (21-38) |
| BMI<sup>a</sup> | 25.2 ± 1.5 (21.6-27.8) | 25.2 ± 1.7 (23.0-28.1) |
| Employment time in the armed forces | $\leq 3$ years: 70 %<br>> 3 years: 30 % | $\leq 3$ years: 64 %<br>> 3 years: 36 % |
| Daily snuff users, $n$ | 9 | 8 |
| Daily smokers, $n$ | 0 | 0 |
| Asthma, allergic rash, pollen or food allergies, $n$ | 9 | 4 |
| Living with pets<sup>b</sup>, $n$ | 2 | 9 |
| Prior regular weekly workouts, $n$ | | |
| < 1 day | 1 | 0 |
| 1-3 days | 1 | 1 |
| 3-5 days | 6 | 9 |
| 5-7 days | 11 | 10 |
| Prior regular use, antiseptic type |  |  |
| ethanol-based | 15 | 20 |
| soap and water (only) | 4 | 0 |
| Antiseptic used during military exercise | ethanol-based: 16<br>none: 1 | chlorine-based: 19 |
| Antiseptic application times during military exercise | Never: 1<br>1-5: 14<br>6-10: 2 | Never: 0<br>1-5: 19<br>6-10: 0 |
| Fraction of time wearing gloves during military exercise, $n$ | | |
| Never: | 0 | 0 |
| 10-40 %: | 0 | 6 |
| 40-80 %:<br>Always: | 11<br>6 | 9<br>4 |
| Total skin samples<sup>c</sup>, *n* | 81 (102) | 82 (110) |
| Hand samples<sup>c</sup>, *n* |  |  |
| Baseline | 19 (19) | 19 (20) |
| Post-intervention | 17 (17) | 19 (19) |
| Follow-up | 15 (15) | 16 (16) |
| Forearm samples<sup>c</sup>, *n* |  |  |
| Baseline | 8 (19) | 5 (20) |
| Post-intervention | 14 (17) | 16 (19) |
| Follow-up | 8 (15) | 7 (16) |

<sup>a</sup>Average values with standard deviations are reported, including the range in brackets.
<sup>b</sup>Living with pets. Control group: 1 with a cat, 1 with a cat and a dog. Intervention group: 7 w/dogs, 2 w/cats.
<sup>c</sup>The number of samples after processing of sequence data (incl. trimming and rarefaction), as well as total collected in parenthesis.

Participants in the intervention group confirmed exclusive use of the chlorine-based antiseptic over the 10-day field exercise. Flask weights showed a median use of 58 mL (range: 14-128 mL), corresponding to an estimated 19 total applications (based on the 3 mL per use), or approximately two applications (range: 1-5) per day. At post-intervention, seven of 19 reported that their hands felt better after using the novel product, while the rest experienced no change in comfort compared to regular practice. Participants in the control group reported using exclusively 85 % ethanol-based disinfection products during the exercise, with the majority (>80 %) reporting one to five applications per day (Table 1). Five out of 16 reported having dry hands after using ethanol-based antiseptic.

From a total of 212 skin swab samples collected and 16S sequenced, 202 samples (106 hands, 96 forearms) were retained after trimming, each with ≥ 5,000 trimmed reads. These yielded 5,537,616 high-quality paired-end reads, with a median of 26,439 reads per sample (range: 2752-74,416). After rarefaction to 7,000 reads, 163 samples (control, 81; intervention, 82) were included in 16S rRNA diversity analyses (Table 1). Rarefaction curves validated this depth as sufficient to capture microbial diversity (Fig. S2). The final dataset comprised 8,536 ASVs, with a median of 329 ASVs per sample (hands: median 394, range 87-932; forearms: median 146, range 44-509).

### Alpha diversity

We investigated temporal changes in alpha diversity on the hands between the chlorine-based intervention group and the control group. A linear mixed model revealed a significant *group* × *time-point* interaction (ANOVA, F = 4.05, *P =* 0.022), indicating differing trajectories of Shannon diversity (Fig. 2). Corresponding results were observed with the Inverse Simpson index (Fig. S3). Initially, we compared baseline to post-intervention changes to assess the immediate effect of antiseptic use. No significant reduction in diversity was found in the intervention group relative to the controls immediately after 10-day use (Shannon estimate: -0.17, *P =* 0.55; Inverse Simpson estimate: -10.2, *P =* 0.45), and antiseptic dosage showed no significant association with post-intervention diversity (effect size Shannon = 0.004, *P =* 0.42; Fig. 2A). However, during the recovery phase (from day 10 to 3W follow-up), the intervention group exhibited a significant increase in diversity compared to the control group (Shannon estimate: +0.82, *P =* 0.0083; Inverse Simpson estimate: +45.8, *P =* 0.0018; Fig 2A, Fig S3, respectively). As the intervention group had more pet owners (Table 1), we tested whether the increase in diversity was correlated with pet ownership and found no association (Pearson’s *r* = 0.096, *P* = 0.72).

**Figure 2:**
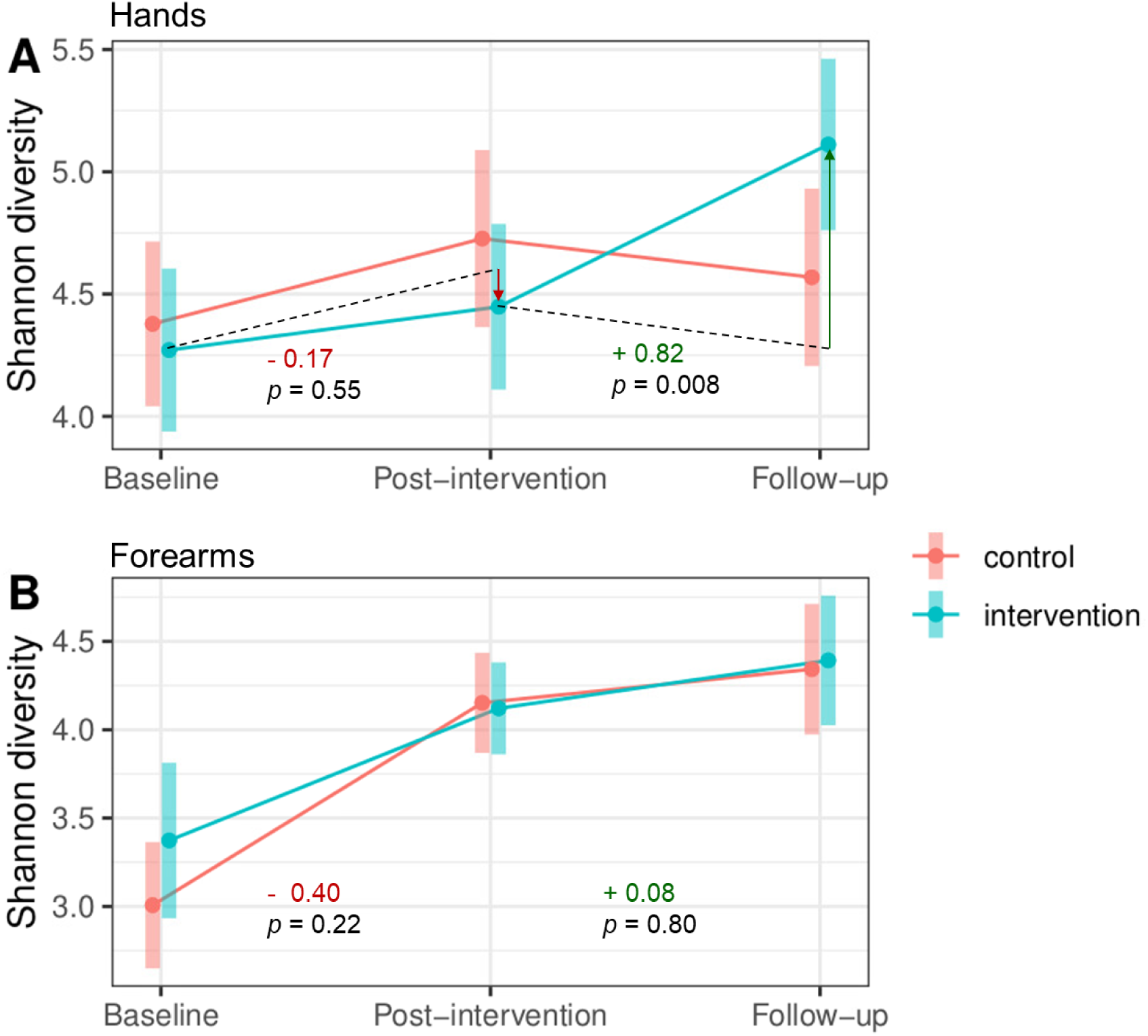
Alpha diversity for group × time point. Interaction plot of Shannon diversity (estimated marginal means) across groups (pink = control, blue = intervention) and time points (baseline, post-intervention: after 10 days of antiseptic use, and follow-up 3 weeks after cessation) on **(A)** hands and **(B)** forearms (untreated control site). The interaction estimate for each time period is annotated along with the *p*-value. Error bars indicate 95 % confidence intervals around the means. Dotted lines in panel (A) demonstrate the counterfactual (where intervention group would be without the intervention), and arrows are added for visualization of the treatment effect.

Forearm samples served as a control site, as the antiseptic was applied only to the hands.

A linear mixed model confirmed that there was no significant interaction between group and time on alpha diversity at this site (F = 0.80, *P =* 0.46; Fig. 2B), with no difference in the change between groups in either time period (*P =* 0.22, *P =* 0.80). Regardless of treatment, both groups exhibited increased forearm diversity during the exercise (*P* < 0.001, Fig. 2B, Fig. S4).

### Beta diversity

To assess whether hand disinfection altered beta diversity beyond the military exercise-induced changes observed in the control group (46), we fitted a PERMANOVA model including a group × time-point interaction. The full model revealed a significant interaction effect across all time points (R^2^ = 0.026, *P* = 0.003), as well as an effect of time point (R^2^ = 0.092, *P* = 0.002), suggesting that temporal trajectories differed between groups, but that the treatment effect was smaller than that of field exercise and recovery (Fig. 3A). When restricting analysis to consecutive two-time point subsets, the interaction effect remained significant during the intervention period (R^2^ = 0.023, *P* = 0.004), but not during the recovery period (R^2^ = 0.014, *P* = 0.17). The untreated forearm site showed no significant group differences over time (R^2^ = 0.028, *P* = 0.44), supporting the hand-specificity of the intervention (Fig. 3C-D). Despite these findings, interpretation of a true treatment effect was complicated by a significant difference between the two groups at baseline (R² = 0.061, *P* = 0.004; Fig. 3A). Groups were also different at post-intervention (*P* = 0.002), but with a similar amount of variation explained (R² = 0.060). At follow-up, the groups showed no significant differences (R² = 0.038, *P* = 0.24). Exploring temporal dynamics within groups, the intervention group exhibited lower variance explained by time point (R^2^ = 0.10) compared to controls (R^2^ = 0.16), suggesting smaller community shifts at the group level. At the individual level, however, there was a tendency for greater changes (intra-individual distances) in the intervention group (Fig. 3B), although this was not statistically significant, neither during the intervention (*P* = 0.3) nor during recovery (*P* = 0.1). Moreover, the amount of antiseptic used by the intervention group was not a significant predictor of beta diversity changes (estimate = 0.023, SE = 0.033, *P =* 0.49).

**Figure 3:**
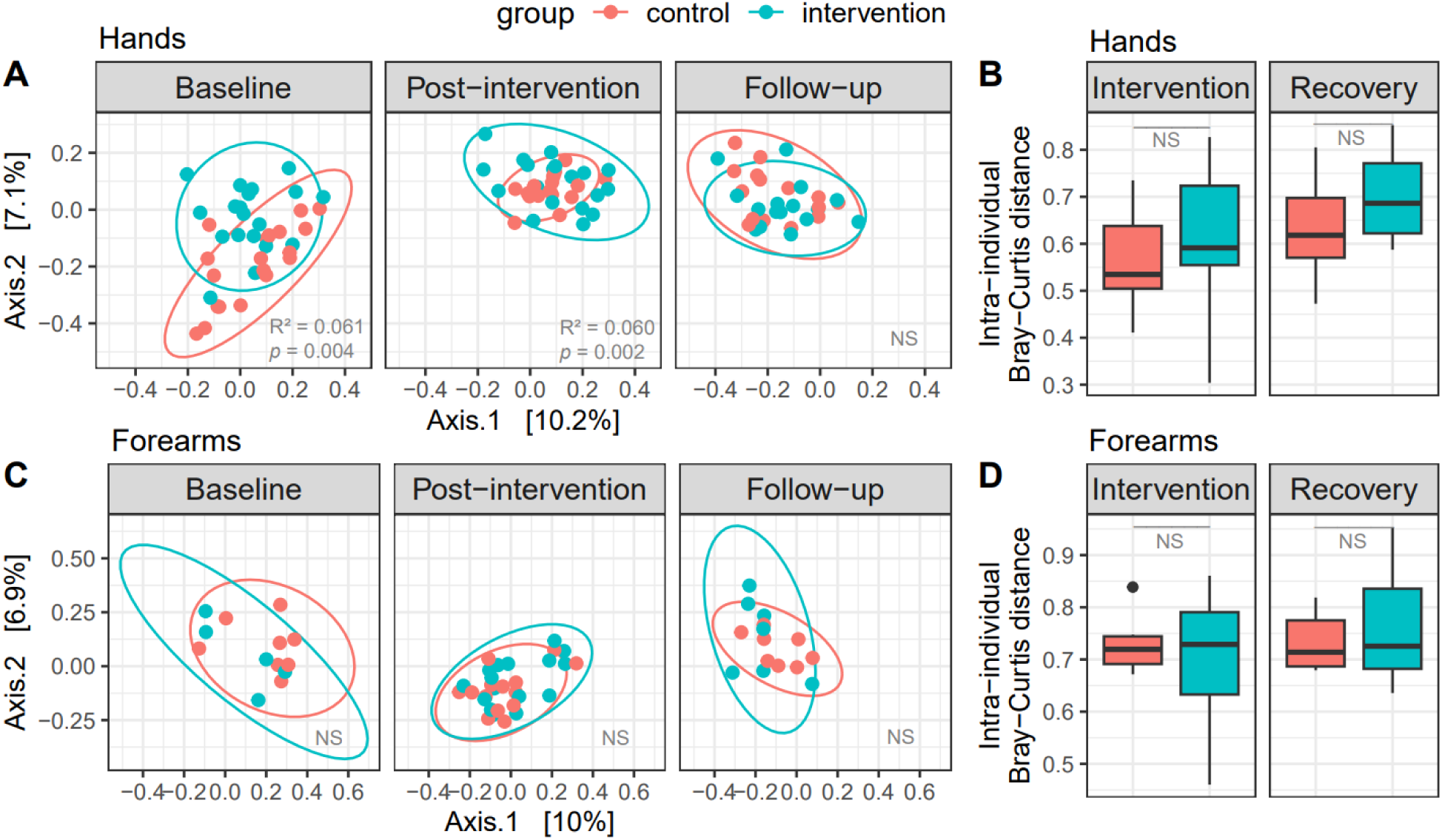
Beta diversity assessed by Bray-Curtis dissimilarity. (**A**, **B**) Hand samples; (**C**, **D**) forearm samples. (**A**, **C**) Principal Coordinates Analysis (PCoA) plots showing group separation (95% CI); points represent samples colored by group (pink = control, blue = intervention). Axes indicate principal components with explained variance (%). Proximity of points indicates high similarity in the microbial community. PERMANOVA: R² is the proportion of variance explained by group differences, with significant differences reported for p < 0.05. NS: non-significant. (**B**, **D**) Box plots of intra-individual distances over the two time periods (intervention: baseline to post-intervention; recovery: post-intervention to follow-up) for each group. Wilcoxon rank-sum test used to compare groups, NS: non-significant.

To assess if group differences in beta diversity were due to changes in variability, we analyzed beta dispersion using PERMDISP. The analysis confirmed overall heterogeneity differences between groups (ANOVA: F = 7.9, *P* < 0.001), with pairwise tests identifying a significant dispersion difference only at post-intervention (permuted *P* = 0.010; Fig. S5A). Accounting for this, ANOSIM confirmed significant group separation at post-intervention (R: 0.12, *P =* 0.003) beyond dispersion effects. Within groups, only the control group exhibited changes in dispersion across time points (ANOVA, F = 13, *P* < 0.001), whereas the intervention group remained comparatively stable (ANOVA, F = 2.8, *P* = 0.07). Both groups appeared more homogeneous at post-intervention (Fig. S5A), although reductions in distances to centroids were not significant (*P* > 0.05). At follow-up, both groups showed similarly elevated dispersion (Fig. S5A). Comparable patterns of beta dispersion were seen at the forearm site (Fig. S5B).

### Differential abundance

Next, we investigated whether particular bacteria were subject to antiseptic intervention-driven abundance shifts. At the chosen threshold (FDR *P* < 0.05, |log2FC| ≥ 1), 18 of 704 evaluated ASVs on hands were significantly enriched or depleted in the intervention group relative to controls in at least one contrast (Fig. 4A). Specifically, we identified nine, seven, and nine ASVs as differentially abundant (DA) between groups for the immediate intervention effect, recovery phase, and the overall study period, respectively, reflecting intervention-driven changes captured by the interaction term. DA ASVs were distributed across three phyla: *Actinomycetota*, *Bacillota*, and *Pseudomonadota*, and all were assigned to distinct genera (Fig. 4A). Seven ASVs appeared as DA in two contrasts, none of which were overlapping for the intervention and recovery period. Clustering of all DA ASVs showed that samples clustered primarily by round, with groups more interspersed (Fig. 4B), consistent with the PERMANOVA results.

**Figure 4:**
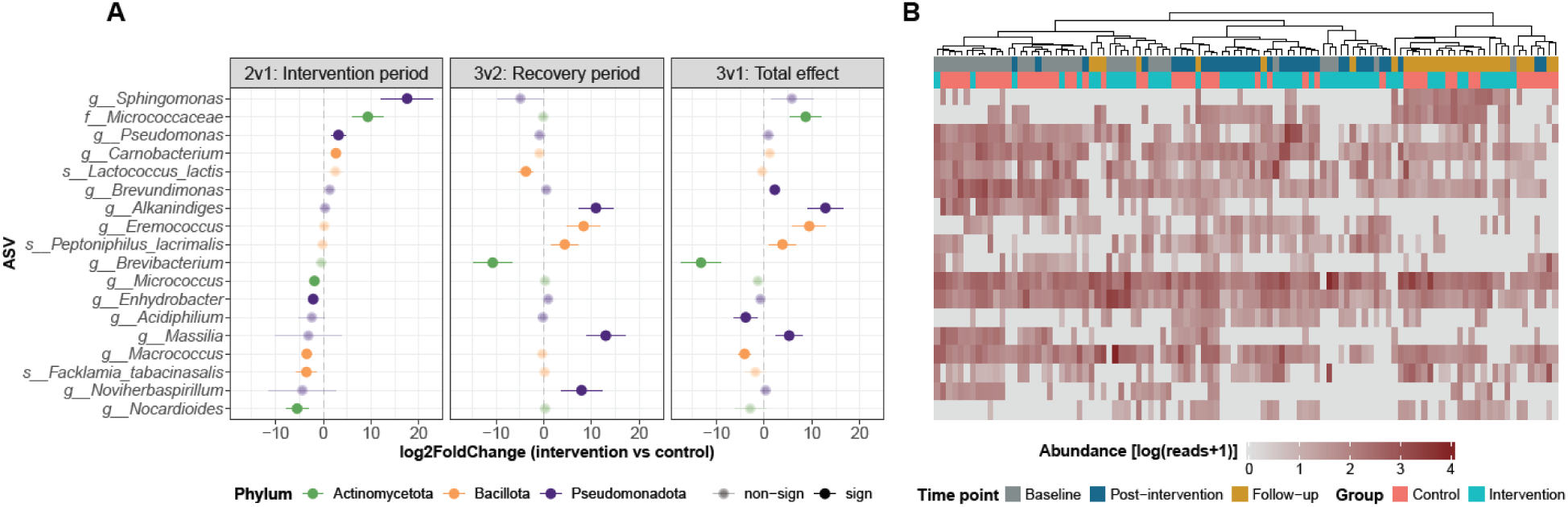
Differentially abundant taxa on hands for group × time. **(A)** ASVs identified by DESeq2 as significantly different between intervention and control groups across three time contrasts (facets). ASVs are colored by phylum and ordered by log2 fold change (± SE) for the intervention period (time point 2 vs. 1). Labels indicate the lowest classified taxonomic rank. Significance thresholds were absolute log2 fold change ≥ 1 and FDR-adjusted *P* < 0.05; opaque points indicate significant ASVs. **(B)** Heatmap of log₁₀(+1)-transformed abundances of differentially abundant ASVs (rows ordered as in panel A) across all hand samples (columns). Samples were hierarchically clustered by Bray–Curtis distances using Ward.D2 linkage. Annotations indicate group (pink = control, blue = intervention) and time point (gray = baseline, blue = post-intervention, gold = follow-up). Color scale ranges from low (gray) to high (red) abundance.

In comparison, in the untreated forearms, there was only one DA ASV for the interaction term out of 312 tested, namely a *Cutibacterium*, which was suppressed in the intervention group during recovery (log2FC = -7.68, *P* < 0.001) and overall (log2FC = -10.4, *P* < 0.001), and was not found to be DA in hands. Complete DESeq2 results are provided in Table S1.

## DISCUSSION

Soldiers are exposed to environmental factors and hygienic challenges during field exercises that change the bacterial communities on their skin (46). In this paper, we reveal subtle compositional shifts during field exercise and a significant increase in diversity three weeks post-treatment when comparing chlorine-based antiseptics to standard hygiene controls, mainly using alcohol-based sanitation. These differences were not seen in untreated forearm samples.

The immediate effects of repeated HOCl-based hand disinfection during the 10-day military field exercise were limited. We found no change in alpha diversity immediately after the exercise, neither within groups nor between groups over time. In contrast, forearm diversity increased similarly in both groups, suggesting a strong environmental effect caused by shared exposure to new microbes in the field (46, 61). It is likely that hand disinfection in both groups suppressed the natural accumulation of environmental microbes, counteracting the diversity increase observed on untreated skin. Vindenes *et al.* recently showed that alpha diversity increases with time since last hand wash (19), supporting this interpretation. However, several studies have also reported minimal short-term effects on overall microbial diversity with normal use of alcohol-based hand sanitizers, indicating a degree of hand microbiome resistance (62, 63). In terms of beta diversity, we found a statistically significant group-by-time interaction, partly reflected by a difference in spread (beta dispersion). It appears that the HOCl-based intervention maintains more individual heterogeneity, not showing the homogenization effect demonstrated in the control group (46). Another study also found greater variation between participants not using alcohol-based hand rubs (63). However, the small effect sizes (R²) of the interaction effects observed in our study indicates that antiseptic type explains only a small fraction of the overall variation. Moreover, baseline differences in composition between groups, likely attributable to pre-existing affiliation with distinct troops, reduced the interpretability of this signal. Only a very small fraction of bacteria (∼2 %) were differentially abundant due to the antiseptic intervention, and with no evident phylogenetic pattern for enriched versus depleted taxa, consistent with the broad-spectrum, non-targeted nature of antiseptics (64).

While the antiseptic intervention did not immediately alter bacterial diversity, the intervention group showed a substantial post-treatment increase relative to controls. This suggests that the non-alcoholic product promoted effects on the skin allowing for higher microbial diversity after cessation of use. By avoiding harsh ingredients, the HOCl-based antiseptic product may preserve conditions that facilitate commensal persistence or rapid recolonization (65, 66). In contrast, the control group’s microbial communities remained suppressed, potentially due to harsher effects of alcohol-based sanitizers, which strip skin lipids that can create a drier environment less hospitable for regrowth (67–70). Follow-up diversity exceeded baseline levels only in the intervention group, which implies a relative resilience of the hand microbiome when not repeatedly exposed to alcohol-based disruption. Several studies agree that the skin microbiome is largely resistant to antiseptics or that perturbation is at least reversible in healthy skin (5, 10, 18, 71); however, these studies primarily examined single applications, which do not correspond to the conditions of frequent use advised in field settings. To our knowledge, the only prior study that has investigated the microbiome response to a hypochlorous acid-based hygiene solution reported no change in bacterial diversity, but this was also only after a single treatment on ocular skin (72). Our findings extend current understanding by demonstrating that formulation-specific effects may be most evident after discontinuation rather than during use. The groups did not differ in their compositional (beta diversity) trajectories during recovery, indicating that the rebound was driven by increases in evenness rather than directional shifts of particular taxa. By follow-up, both groups show similar, elevated within-group dispersion, reflecting a reassembly phase where host factors dominate. As the soldiers were on military leave during the three week recovery period and thus no longer belonged to any group/troop, this loss of group signature and convergence toward high individualized variability was as expected. Our results indicate that while the difference in hygienic practices was overshadowed by environmental pressure during use, the non-alcoholic formulation was associated with greater recolonization potential once both perturbations ceased. Although this study cannot directly assess pathogenicity or confirm that higher diversity is beneficial, prior research links lower hand microbial diversity to increased presence of potential pathogens (63). Based on this, and its potential for improved wound healing and reduced subjective discomfort (29, 32, 34, 39, 43), the chlorine-based antiseptic is a promising alternative to alcohol-based products for military personnel.

A key strength of this study is its interaction design, which enabled us to assess differences in microbiome trajectories between groups over time. This approach, combined with longitudinal sampling at three time points, allowed us to capture both immediate and sustained microbiome responses to repeated antiseptic intervention - dynamics often missed in studies limited to single-dose or short-term assessments (5, 10, 12, 18). The inclusion of an untreated control site (forearms) for each participant, which is rare in similar studies (10, 65), served as an internal reference strengthening interpretation of antiseptic-driven changes. Nonetheless, our study has several limitations. Firstly, full randomization was not feasible due to logistical constraints, resulting in baseline differences in beta diversity between groups. However, the lack of group differences in forearm microbiota suggests that observed hand-specific effects likely stem from the antiseptic intervention rather than troop-related factors. Furthermore, antiseptic usage was relatively low and variable across participants, with potential for minor product loss during application. Still, self-reported application frequencies aligned with measured quantities and were similar across both groups. While dose-response analyses should be interpreted with caution, they suggest that dosage had minimal impact on the primary outcomes. Finally, while not all contributing factors could be controlled, there is a strength in evaluating the intervention under operationally relevant conditions. Future research should investigate the effects of prolonged antiseptic use, tracking soldiers over extended periods to correlate microbiome changes and antiseptics with reported illness, leveraging their monitored and uniform lifestyle.

In summary, our research suggests that environmental context exerts a stronger influence on skin microbial community structure than repeated chlorine-based antiseptic use, particularly under high-exposure conditions. The impact of antiseptic use may manifest most strongly after product discontinuation rather than during treatment, and chlorine-based formulations may better support re-establishment of a diverse microbiome compared to standard alcohol-based practices.

## DATA AVAILABILITY STATEMENT

The sequencing data have been deposited in the NCBI Short Read Archive under BioProject accession <u>PRJNA1159222</u>. Code used to analyze the data can be found at: https://github.com/Rounge-lab/skin_microbiome_16S_seq_analyses.

## CONFLICT OF INTEREST

The authors state no conflict of interest.

## ACKNOWLEDGEMENTS

This study was funded by the Norwegian Research Council through an industrial PhD grant to SoftOx Solutions AS (project number 332591). We acknowledge Clinical Microbiomics AS (Copenhagen, Denmark) for performing high-throughput DNA extractions of all skin samples, and the Norwegian Sequencing Centre (Oslo, Norway) for library preparation and sequencing. Data analysis was performed on resources from Sigma2 - the National Infrastructure for

High-Performance Computing and Data Storage in Norway. We also declare the assistance of OpenAI’s ChatGPT (versions 4.0 and 5.0) in providing programming syntax and linguistic suggestions, while the authors take full responsibility for the content of the paper.

## AUTHOR CONTRIBUTIONS

Conceptualization: SG, ØAV, TBR, OAØ, RJSO; Funding Acquisition: SG, ØAV, OAØ; Project Administration: SG, ØAV, OAØ; Investigation: SG, ØAV, MD, GDG; Resources: MD, GDG; Data Curation: SG; Formal Analysis: SG, EEB; Software: SG, EEB; Visualization: SG, EEB; Supervision: OAØ, TBR, RJSO; Writing - Original Draft Preparation: SG; Writing - Review and Editing: SG, EEB, OAØ, TBR, RJSO, ØAV, MD, GDG.

## ETHICS STATEMENT

The study and experimental procedures were reviewed and approved by the Regional Committees for Medical and Health Research Ethics in South-Eastern Norway (ref: 359040, approved 03-02-2022). All participants provided their written informed consent to participate in this study.

## SUPPLEMENTAL MATERIAL

### Supplementary figures

**Figure S1:**
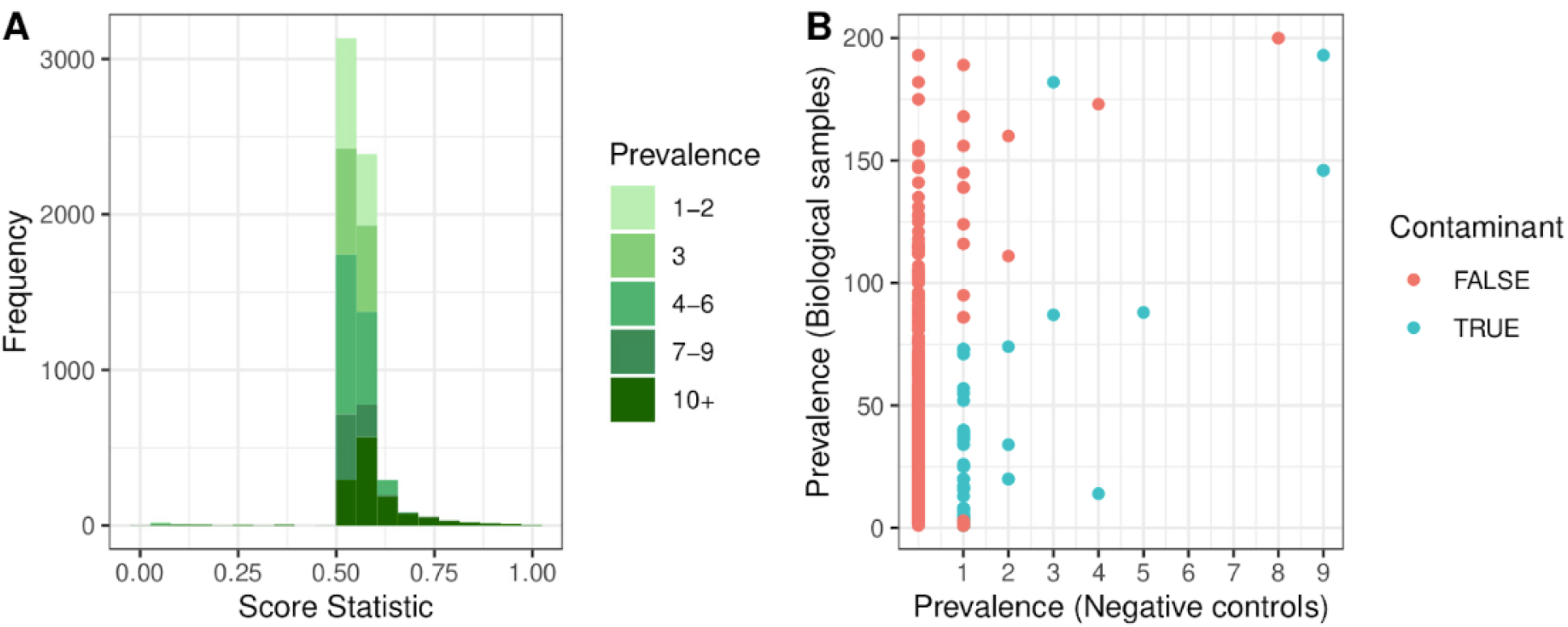
***Decontam()* results. (A)** Distribution of score statistics from *decontam* assigned to each amplicon sequence variant (ASV) based on prevalence (color intensity indicating the total number of samples each ASV was present in (i.e, prevalence). A score < 0.5 indicates a contaminant. **(B)** Prevalence plot of present/absent ASVs in true samples vs negative controls. 52 contaminant ASVs identified by *decontam* (with prevalence method) are colored blue. These represented 440,796 reads, accounting for ∼9% of the (non-chimeric) reads overall.

**Figure S2:**
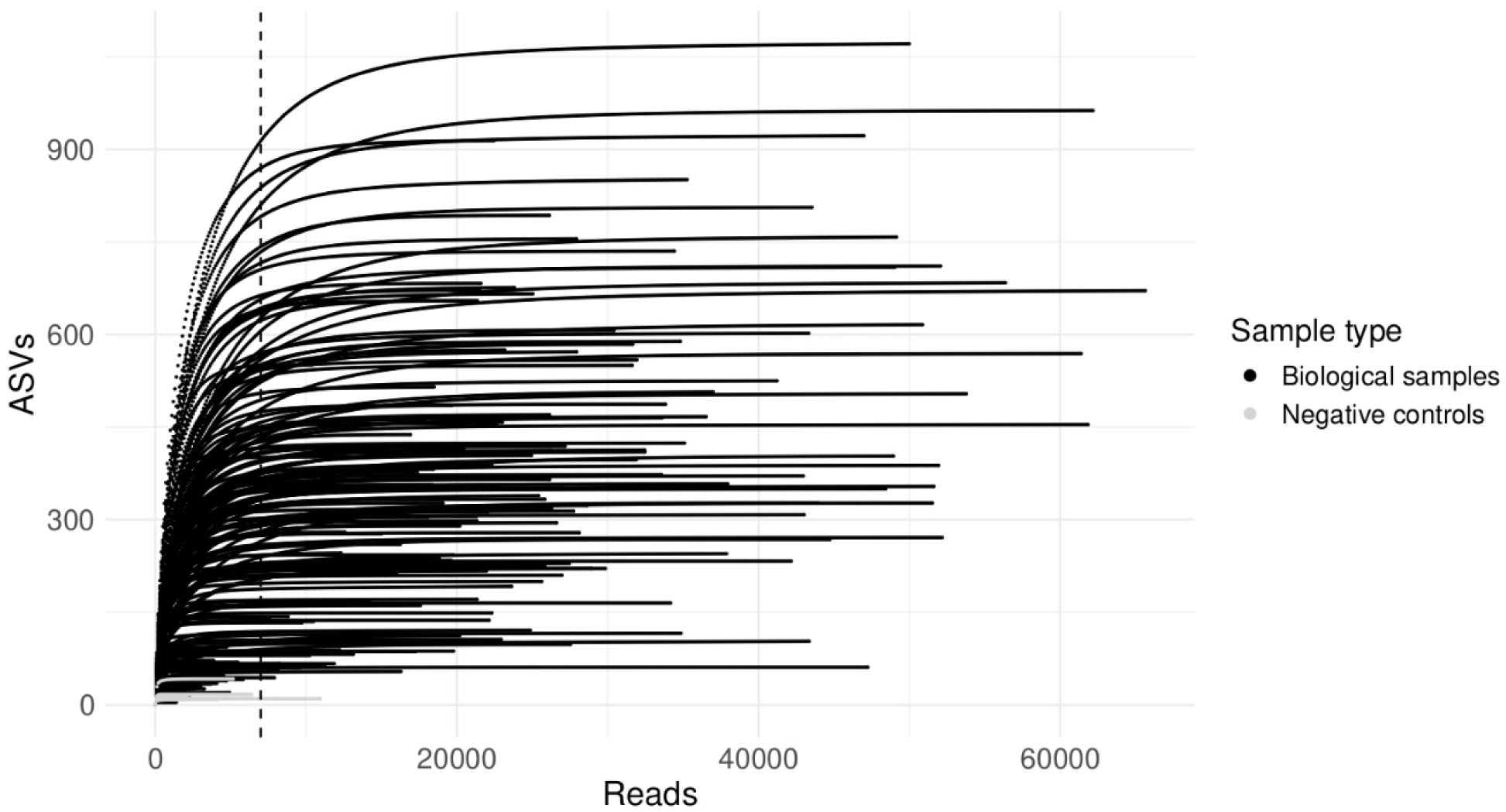
Rarefaction curves. Number of amplicon sequence variants (ASVs) per sample size (number of trimmed reads) for all biological samples (black, n=202) and negative controls (gray, n=9). The dotted line represents the chosen sampling depth of 7000 reads, which retains most of the data’s diversity (where most samples have reached a plateau of new ASVs per increased depth) and the number of samples.

**Figure S3:**
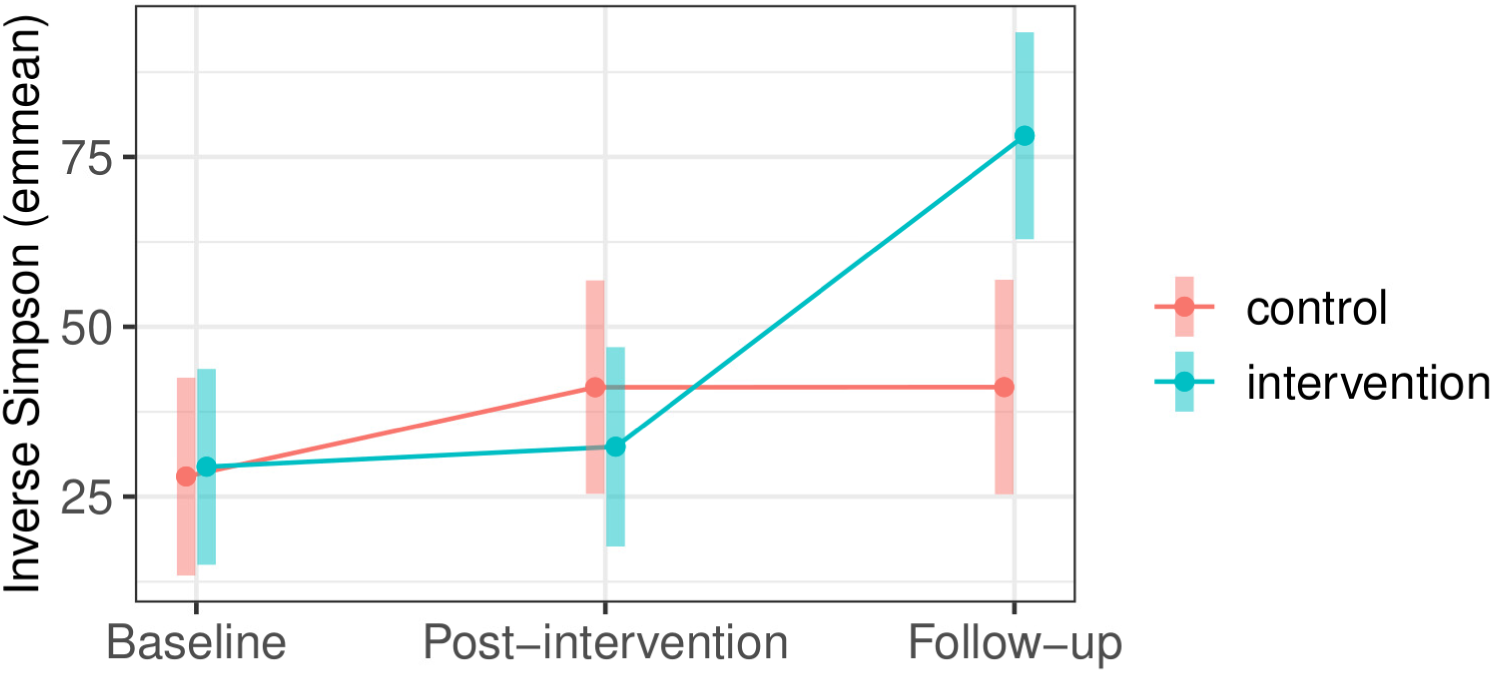
Inverse Simpson. Estimated marginal means of Inverse Simpson diversity for hand samples over time, comparing the two groups. This metric exhibits a similar temporal pattern to Shannon diversity (see Fig. 1A).

**Figure S4:**
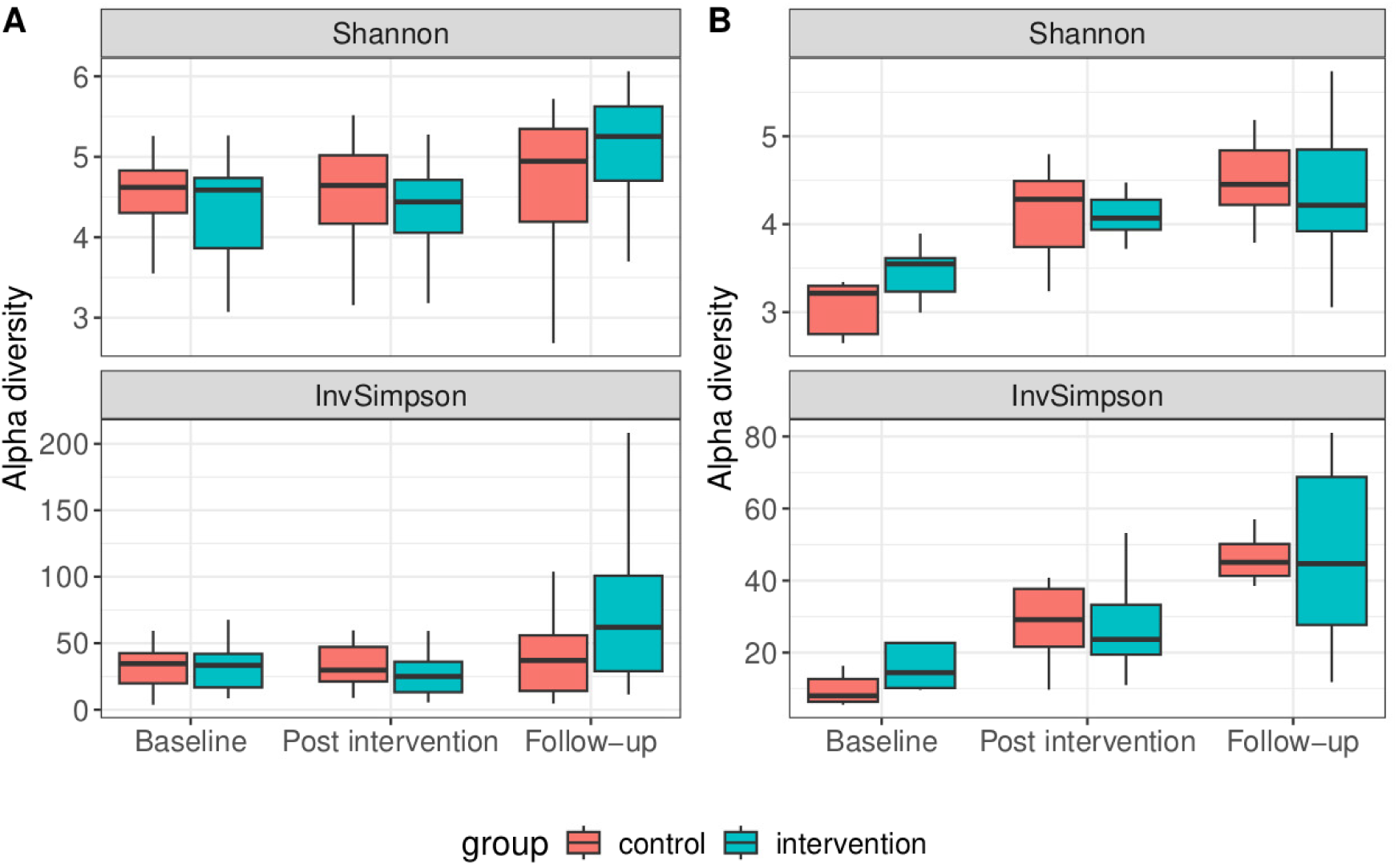
Alpha diversity (unadjusted) over time. (**A**) hands, (**B**) forearms. (Unadjusted) Shannon and Inverse Simpson diversity metrics for each group (pink = control, blue = intervention) over time points.

**Figure S5:**
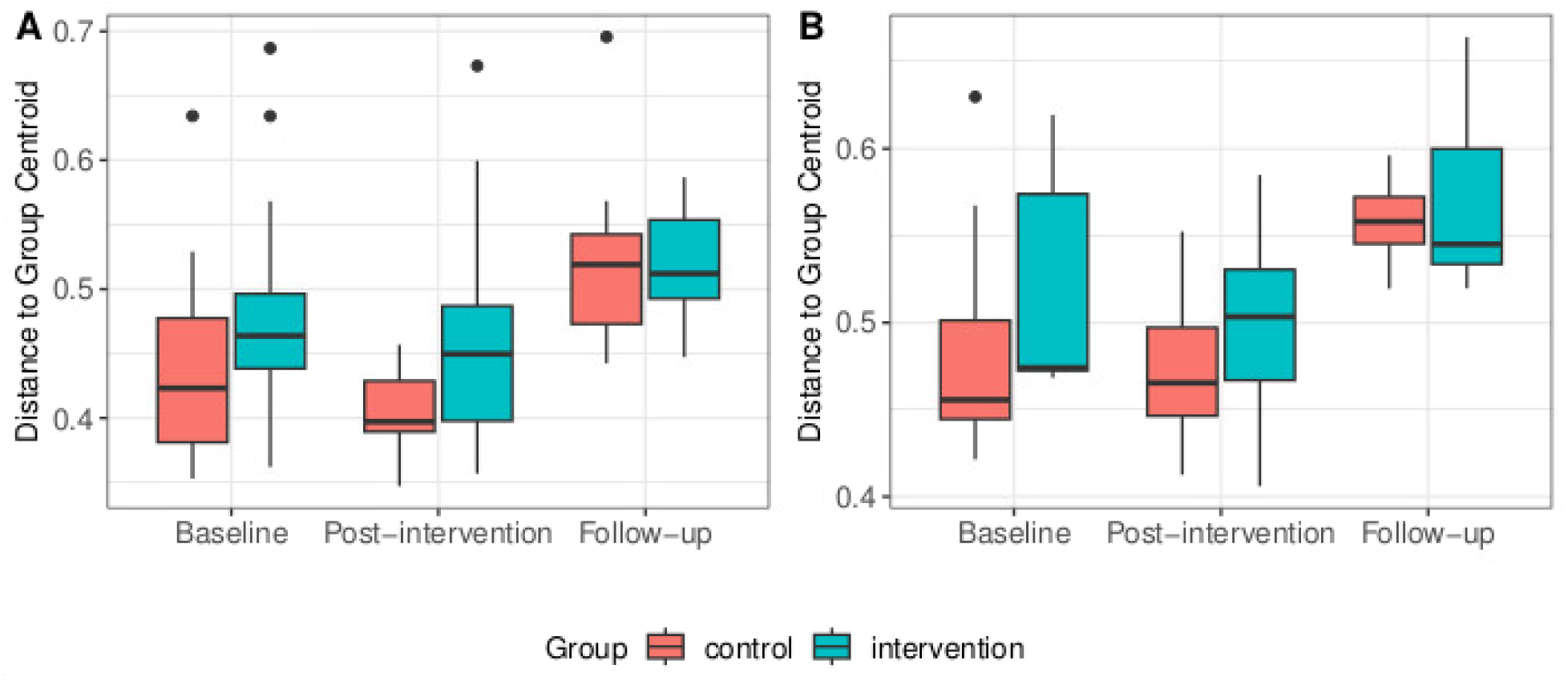
**Beta dispersion**. Distances to group centroids for each group (pink = control, blue = intervention) and time point in each skin site, **(A)** hands, **(B)** forearms.

### Supplementary tables

Table S1: Differential abundance table (uploaded)

## Notes

### Competing Interest Statement

The authors have declared no competing interest.

