## Supplementary information for "Longitudinal analysis of the hand microbiome in response to chlorine-based antiseptic use during a military field exercise"

### SUPPLEMENTAL MATERIAL

#### Supplementary figures

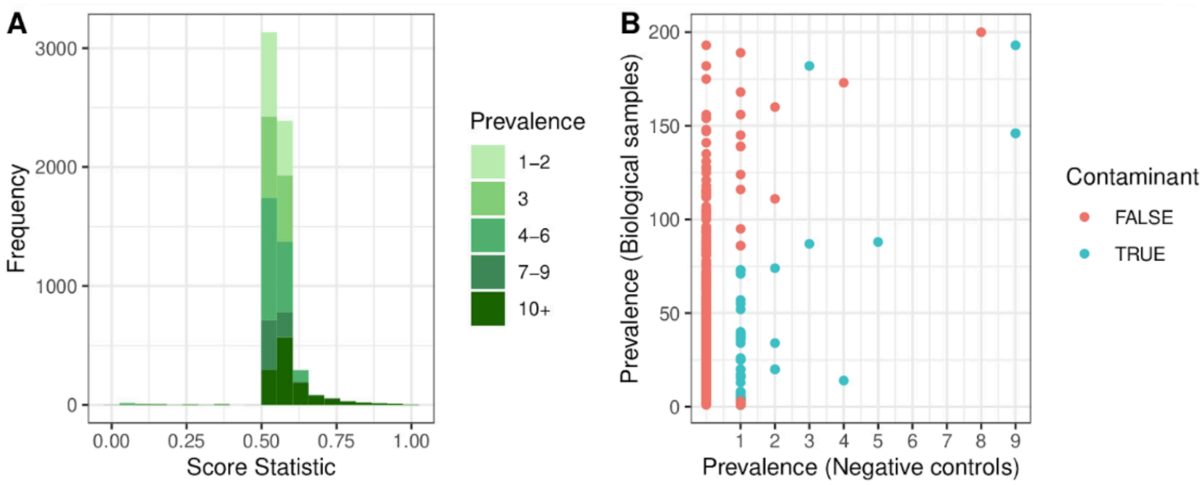


**Figure S1**: ***Decontam()* results. (A)** Distribution of score statistics from *decontam* assigned to each amplicon sequence variant (ASV) based on prevalence (color intensity indicating the total number of samples each ASV was present in (i.e, prevalence). A score < 0.5 indicates a contaminant. **(B)** Prevalence plot of present/absent ASVs in true samples vs negative controls. 52 contaminant ASVs identified by *decontam* (with prevalence method) are colored blue. These represented 440,796 reads, accounting for ~9% of the (non-chimeric) reads overall.

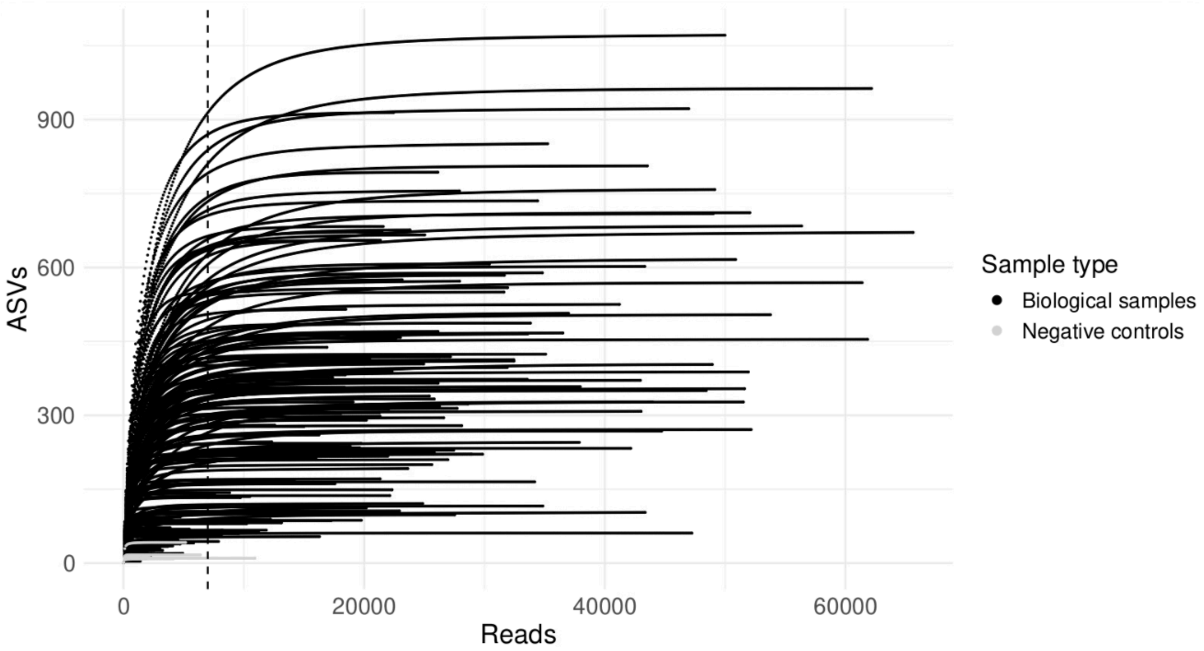


**Figure S2**: **Rarefaction curves.** Number of amplicon sequence variants (ASVs) per sample size (number of trimmed reads) for all biological samples (black, n=202) and negative controls (gray, n=9). The dotted line represents the chosen sampling depth of 7000 reads, which retains most of the data's diversity (where most samples have reached a plateau of new ASVs per increased depth) and the number of samples.

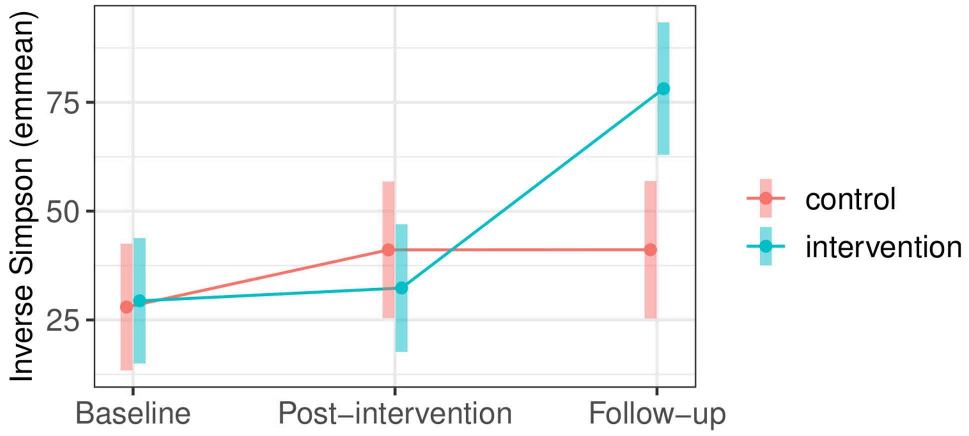


**Figure S3: Inverse Simpson.** Estimated marginal means of Inverse Simpson diversity for hand samples over time, comparing the two groups. This metric exhibits a similar temporal pattern to Shannon diversity (see Fig. 1A).

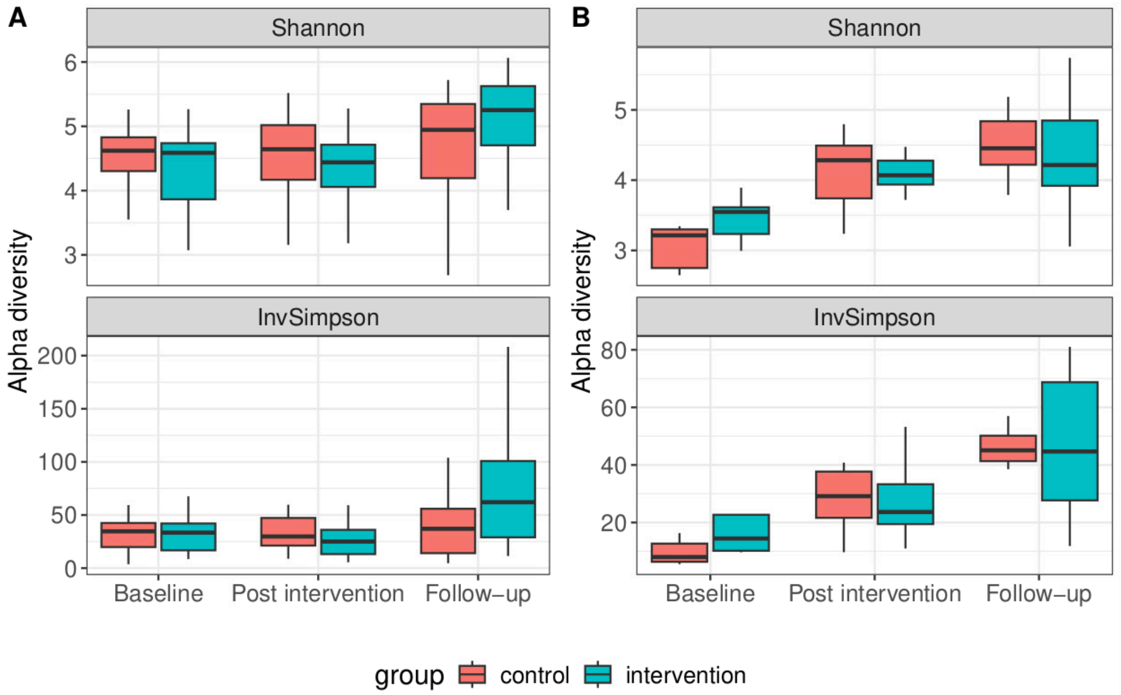


**Figure S4**: **Alpha diversity (unadjusted) over time.** (**A**) hands, (**B**) forearms. (Unadjusted) Shannon and Inverse Simpson diversity metrics for each group (pink = control, blue = intervention) over time points.

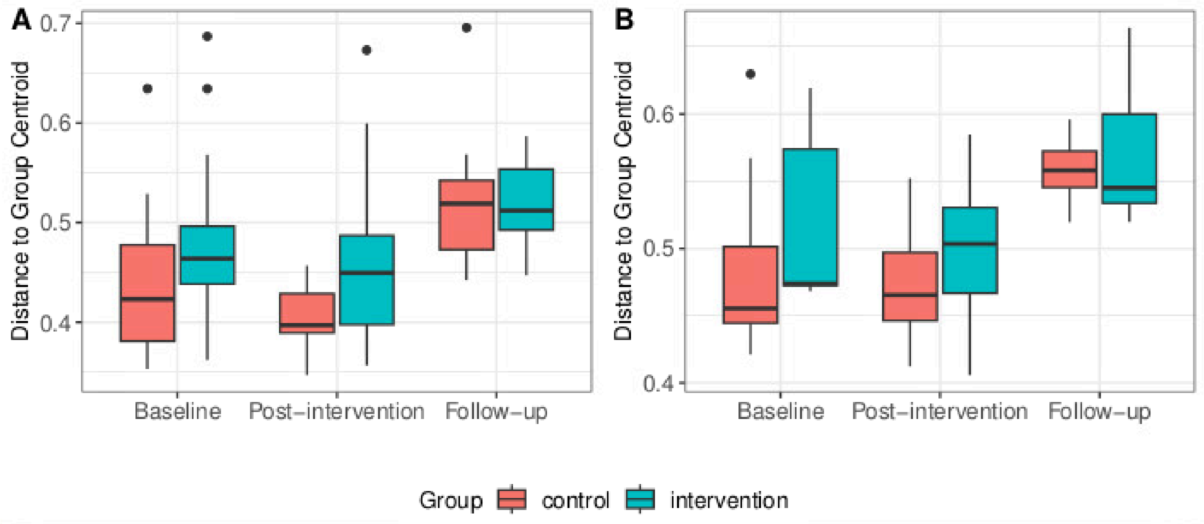


**Figure S5**: **Beta dispersion**. Distances to group centroids for each group (pink = control, blue = intervention) and time point in each skin site, **(A)** hands, **(B)** forearms.

#### Supplementary tables

**Table S1**: Differential abundance table (uploaded)
